# Machine Learning Based Refined Differential Gene Expression Analysis of Pediatric Sepsis

**DOI:** 10.1101/2020.02.21.959494

**Authors:** Mostafa Abbas, Yasser EL-Manzalawy

## Abstract

**Background:** Differential expression (DE) analysis of transcriptomic data enables genome-wide analysis of gene expression changes associated with biological conditions of interest. Such analysis often provide a wide list of genes that are differentially expressed between two or more groups. In general, identified differentially expressed genes (DEGs) can be subject to further downstream analysis for obtaining more biological insights such as determining enriched functional pathways or gene ontologies. Furthermore, DEGs are treated as candidate biomarkers and a small set of DEGs might be identified as biomarkers using either biological knowledge or data-driven approaches.

**Methods:** In this work, we present a novel approach for identifying biomarkers from a list of DEGs by re-ranking them according to the Minimum Redundancy Maximum Relevance (MRMR) criteria using repeated cross-validation feature selection procedure.

**Results:** Using gene expression profiles for 199 children with sepsis and septic shock, we identify 108 DEGs and propose a 10-gene signature for reliably predicting pediatric sepsis mortality with an estimated Area Under ROC (AUC) score of 0.89.

**Conclusions:** Machine learning based refinement of DE analysis is a promising tool for prioritizing DEGs and discovering biomarkers from gene expression profiles. Moreover, our reported 10-gene signature for pediatric sepsis mortality may facilitate the development of reliable diagnosis and prognosis biomarkers for sepsis.

## Background

Pediatric sepsis is a life-threatening condition that is considered a leading cause of morbidity and mortality in infants and children [1, 2]. Sepsis is a systematic response to infection that is characterized by a generalised pro-inflammatory cascade, which may lead to extensive tissue damage [3]. Early recognition of sepsis and septic shock will help pediatric care physicians to intervene before the onset of advanced organ dysfunction and thus reduce the mortality and length of stay as well as post critical care complications [4]. However, reliable risk stratification of sepsis, especially in children, is a challenge due to significant patient heterogeneity [5] and existing poor definitions of sepsis in pediatric populations [6].

Existing physiological scoring tools in ICU, such as Acute Physiologic and Chronic Health Evaluation (APACHE) [7] and Sepsis-related Organ Failure Assessment (SOFA) [8], use clinical and laboratory measurements to quantify critical illness severity but provide little information about the risk for poor outcome (e.g., mortality) at the onset of the disease [2]. Several recent studies have proposed sepsis prognostic biomarkers (e.g., [9, 5, 10]) as well as sepsis diagnostic biomarkers (e.g., [11, 12, 13]) by differentiating between infectious and non-infectious systemic inflammatory response syndrome. To date, transcriptomic, proteomic, and metabolomic data have been used to identify sets of genes, proteins, or metabolites that are differentially expressed among patients [14]. However, a major challenge for developing clinically feasible sepsis biomarkers is to have a fast turnaround time [14, 15].

Recent advances in high-throughput transcriptomic technology have created opportunities for precision critical care medicine by enabling fast and clinically feasible profiling of gene expressions within few hours. For example, Wong et al. [16] used a multiplex messenger RNA quantification platform (NanoString nCounter) to profile the expressions of previously identified 100 three subclass-defining genes [17] in 8-12 hours. Differential gene expression (DGE) analysis is a commonly used computational approach for identifying genes whose expressions are significantly different between two phenotypes. Given gene expression profiles for septic patients annotated with targeted outcome (e.g., survival vs. non-survival), DGE analysis typically associates a *p*-value (that could be corrected for multiple hypothesis testing) with each gene from the two groups (e.g. survivals and non-survivals). Then, DEGs are those genes with *p*-values lower than a specific threshold (typically, 0.05) and user-specified thresholds for fold change (FC) for up- and down-regulated genes [18]. A typical DE analysis of gene expression profiles returns hundred or more DEGs, where considerable number of them might be highly correlated with one or more other DEGs.

Against this background, we present a novel method for refining the results of the statistical DE analysis methods via re-ranking and prioritizing the genes from the outcome of DE analysis. Specifically, we propose a hybrid approach that leverages: i) statistical DE analysis for identifying a wide list of DEGs; ii) supervised feature selection methods for selecting an optimal subset of DEGs with maximum relevance for predicting the target variable and minimum redundancy among selected genes; iii) supervised machine learning methods for assessing the discriminatory power of the selected genes. Using gene expression profiles from the blood samples extracted from 199 children admitted to ICU and diagnosed with sepsis or septic shock, we first report a list of 108 DEGs and associated enriched functional pathways. Then, we demonstrate the viability of our proposed gene re-ranking methods in identifying a 10-gene signature for mortality in pediatric sepsis.

## Methods

### Data

Normalized and pre-processed transcriptomic gene expression profiles were downloaded from [19]. These gene expression profiles represent peripheral blood samples collected from 199 pediatric patients (later diagnosed with sepsis or septic shock) during the first 24 hours of admission to the pediatric ICU. Affymetrix CEL files were downloaded from NCBI GEO accession number GSE66099 and re-normalized using the gcRMA method in affy R package [20]. Probe-to-gene mappings were downloaded from the most recent SOFT files in GEO and the mean of the probes for common genes were set as the gene expression level.

### Differential expression analysis

We used limma R package (Version 3.42.0) [18] to identify the deferentially expressed genes with a Benjamini-Hochberg (BH) correction method. We calculated the fold change with respect to the non-survival (i.e., the upregulated genes are the genes with expression of the non-survival samples that are higher than the expression of these genes in the survival samples).

### Classification methods

We experimented with three commonly used machine learning algorithms for developing and evaluated binary classifiers for predicting mortality in pediatric sepsis: i) Random Forest [21] with 100 trees (RF100); ii) eXtreme Gradient Boosting [22] with 100 weak tree learners (XGB100); iii) Logistic Regression (LR) [23] with L2 regularization. The three algorithms are implemented in the Scikit-learn machine learning library [24].

### Feature selection methods

We used two feature selection methods that have been widely used with gene expression data, Random Forest Feature Importance (RFFI) [21] and Minimum Redundancy and Maximum Relevance (MRMR) [25]. For the RFFI method, we trained a RF with 100 trees and then feature importance scores which quantify the contribution of each feature in the learned RF model were used to sort and rank the input features and only top *k* = 1, 2, …, 10 were selected for training our classifiers. For MRMR feature selection method, we used the training data to select the top *k* features. These feature were selected such that the objective function in Eq. 1 is maximized. Let, Ω, *S*, and Ω_*S*_ denote input, selected, and non-selected input features, respectively. The first term in Eq. 1 uses a relevance function *f* (*x*_*i*_, *y*) to quantify the relevance of the feature *x*_*i*_ for predicting the target output *y* while the second term quantifies the redundancy among the selected features in *S* using the function *g*(*x*_*j*_, *x*_*l*_). We implemented the MRMR algorithm [25, 26] as a Scikit-learn feature selection model using Python. In our experiments, we use the Scipy implementation of the Pearson correlation coefficient to compute redundancy between features and for relevance we tried three functions (implemented in scikit-learn): area under ROC curve (MRMR_auc); *χ*^2^ (MRMR_chi2); and F-Statistic (MRMR_fstat).

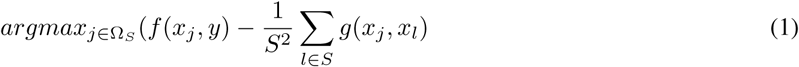

### Marker genes discovery and performance evaluation

We identified top discriminative features (i.e., marker genes) and estimated the performance of the machine learning classifiers using 10 runs of the 10-fold cross-validation procedure. Briefly, we repeated the following procedure 10 times: First, the dataset was randomly partitioned into 10 equal subsets (each with the same survival to non-survival ratio as the entire dataset). Nine of the 10 subsets were combined to serve as the feature selection and training set while the remaining subset was held out for estimating the performance of the trained classifier. This procedure was repeated 10 times, by setting aside a different subset of the data as the test set. Overall, we had 100 runs of train and test experiments. The reported performance is the the average of the 100 runs of performance estimates using the test sets and the score of each feature represents the fraction of how many times this feature was selected in the 100 runs (i.e., a feature with a score of 0.85 means that this feature had been selected to train the classifier in 85 out of 100 runs).

We assessed the performance of classifiers using five widely used predictive performance metrics [27]: accuracy (ACC), Sensitivity (Sn); Specificity (Sp); and Matthews correlation coefficient (MCC); Area under ROC curve (AUC) [28]. AUC is a widely used metric and summary statistic of the ROC curve. However, when several models have almost the same AUC score, we can still compare them by examining their ROC curves to determine if a model has an ROC curve that completely or partially (in the leftmost region) dominates all other ROC curves.

### Pathway enrichment analysis

We used the function *find_enriched_pathway* in the KEGGprofile R package (Version 1.28.0) [29] to map the deferentially expressed genes in KEGG pathway database [30]. In our experiments, pathways with adjusted *p*-value ≤ 0.05 and gene count ≥ 2 were considered significantly enriched.

## Results

### Identification of differentially expressed genes and enriched pathways

Based on absolute fold change ≥ 1.5 and adjusted *p*-value ≤ 0.05, 108 from a total of 10,596 genes were found to be DEGs between survival and non-survival septic pediatric patients (See Additional file 1: Table S1) and Additional file 2: Figure S1). Table 1 shows the top 10 DEGs when the genes are ranked using the absolute value of their fold change. Only one gene, TGFBI, is down-regulated while the remaining nine genes are up-regulated. TGFBI is among the 11 genes that have been used in the Sepsis MetaScore (SMS) gene expression diagnostic method [11, 31]. The top three upregulated genes are SLC39A8, RHAG, and DDIT4. SLC39A8 is found in the plasma membrane and mitochondria and plays a critical role at the onset of inflammation [32]. Both RHAG (also called SLC42A1) and SLC39A8 belong to solute carrier (SLC) group of membrane transport proteins. Finally, increased expressions of DNA Damage Inducible Transcript 4 (DDIT4) gene had been associated with higher risks of mortality in sepsis patients [19, 10].

**Table 1:**
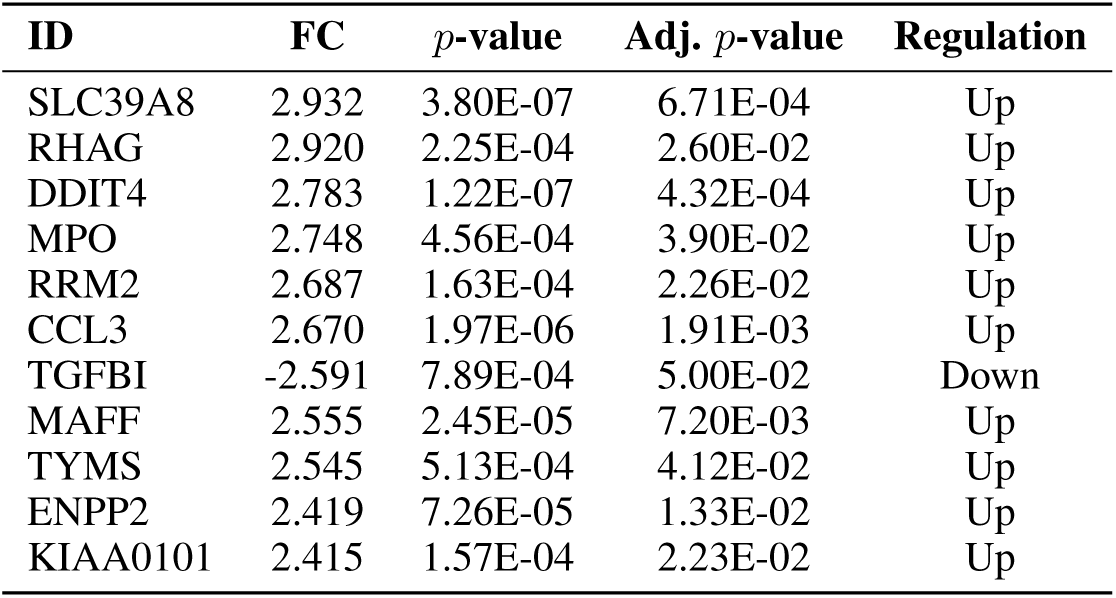
List of top 10 DEGs ranked by the absolute value of the fold change.

In order to get biological insights into the functional rules of the identified 108 DEGs, we used the KEGGProfile R package to identify enriched human KEGG pathways in this set of genes. In our experiments, we did not threshold on the *p*-value, adjusted *p*-value, or minimum number of genes in the pathway such that the returned results include all KEGG pathways that have at least one gene in common with the target set of genes. The complete set of results is provided in Additional file 1: Table S2. We considered a pathway to be significantly enriched if its adjusted *p*-value is ≤ 0.05 and at least two DEGs are included in that pathway. Using this criteria, we got 8 significantly enriched pathways (Table 2). Most of these pathways had been linked to inflammation and/or DNA damage.

**Table 2:** List of significantly enriched KEGG pathways.

| Pathway | *p*-value | Adj. *p*-value |
| --- | --- | --- |
| Cell cycle | 5.71E-12 | 1.92E-09 |
| DNA replication | 8.02E-09 | 1.35E-06 |
| Oocyte meiosis | 5.02E-06 | 4.23E-04 |
| Mineral absorption | 4.78E-06 | 4.23E-04 |
| p53 signaling pathway | 2.13E-04 | 1.23E-02 |
| Human T-cell leukemia virus 1 infection | 2.18E-04 | 1.23E-02 |
| Pyrimidine metabolism | 9.22E-04 | 3.89E-02 |
| Progesterone-mediated oocyte maturation | 9.16E-04 | 3.89E-02 |

Additional file 2: Figure S2 shows the heatmap of the correlation matrix of the 108 DEGs. The figure shows that up-regulated and down-regulated DEGs are clustered separately. We also noted that within each cluster, every gene might be highly correlated with multiple other genes.

### Can a small subset of the DEGs discriminate between survivals and non-survivals?

Here, we report the results of evaluating 120 models obtained using a combination of three supervised classification algorithms, four feature selection methods, and 10 possible values for the number of selected features (*k* = {1, 2, …, 10}). Additional file 1: Table S3 shows the average performance metrics estimated over 10 runs of 10-fold cross-validation experiments. Figure 1 shows the boxplots of the average AUC scores for each combination of a classification algorithm and a feature selection method. Interestingly, MRMR_auc is consistently the best feature selection method using any of the three classification algorithms considered in our experiments. Surprisingly, we found that the models obtained using this feature selection method and LR algorithm not only have the best performance (in terms of AUC scores) but also have the lowest variance in estimated AUC (i.e., AUC scores are between 0.84 and 0.85). Figure 2 shows that (using MRMR_auc feature selection) LR models outperformed corresponding RF100 and XGB100 models for any choice of the number of selected features in *k* = {1, 2, …, 10} Using this figure, one might conclude that the best model with AUC score of 0.85 is using only 2 selected features. However, to accurately identify the best performing LR model, we inspected the average ROC curves of these LR models (See Additional file 2: Figure S3). The LR model using only 2 features is dominated in the leftmost region of the curve (i.e., region corresponds to specificity greater than 0.80) by all other models. For a target specificity greater than 0.80, the best ROC curve corresponds to the model trained using top seven selected DEGs. We concluded that the best model (out of the 120 models evaluated in this study) is based on LR algorithm and MRMR_auc method for selecting top seven DEGs. Therefore, only seven genes are needed to achieve the highest AUC score of 0.85.

**Fig. 1.**
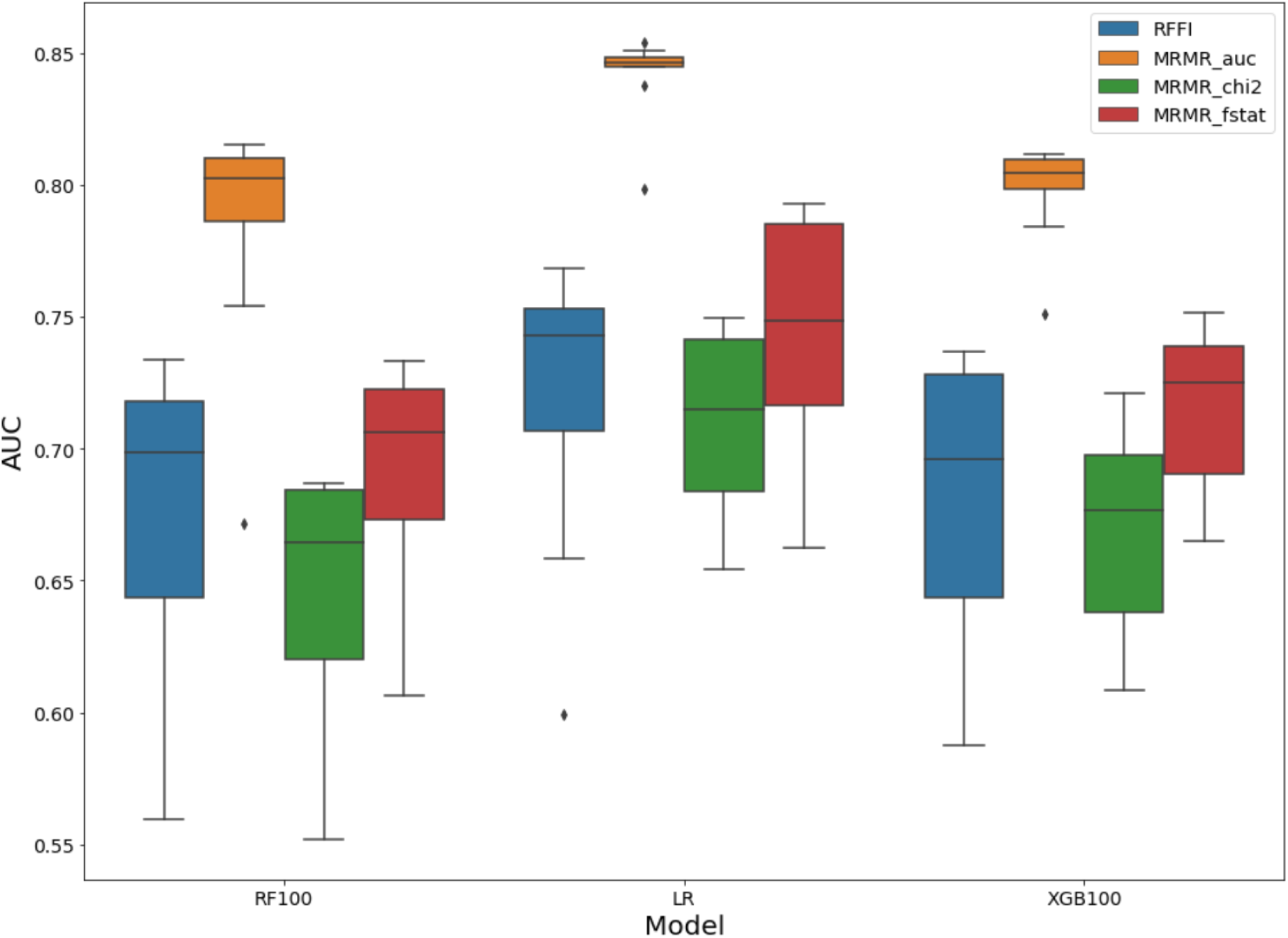
Comparisons of (a) LR (b) RF100 and (c) XGB100 classifiers evaluated using four different feature selection methods and 10 runs of 10-fold cross-validation experiments. Each boxplot represents the distribution of average AUC score of 10 models evaluated using a given classification algorithm and feature selection method for selecting top *k* = 1, 2, …, 10 features.

**Fig. 2.**
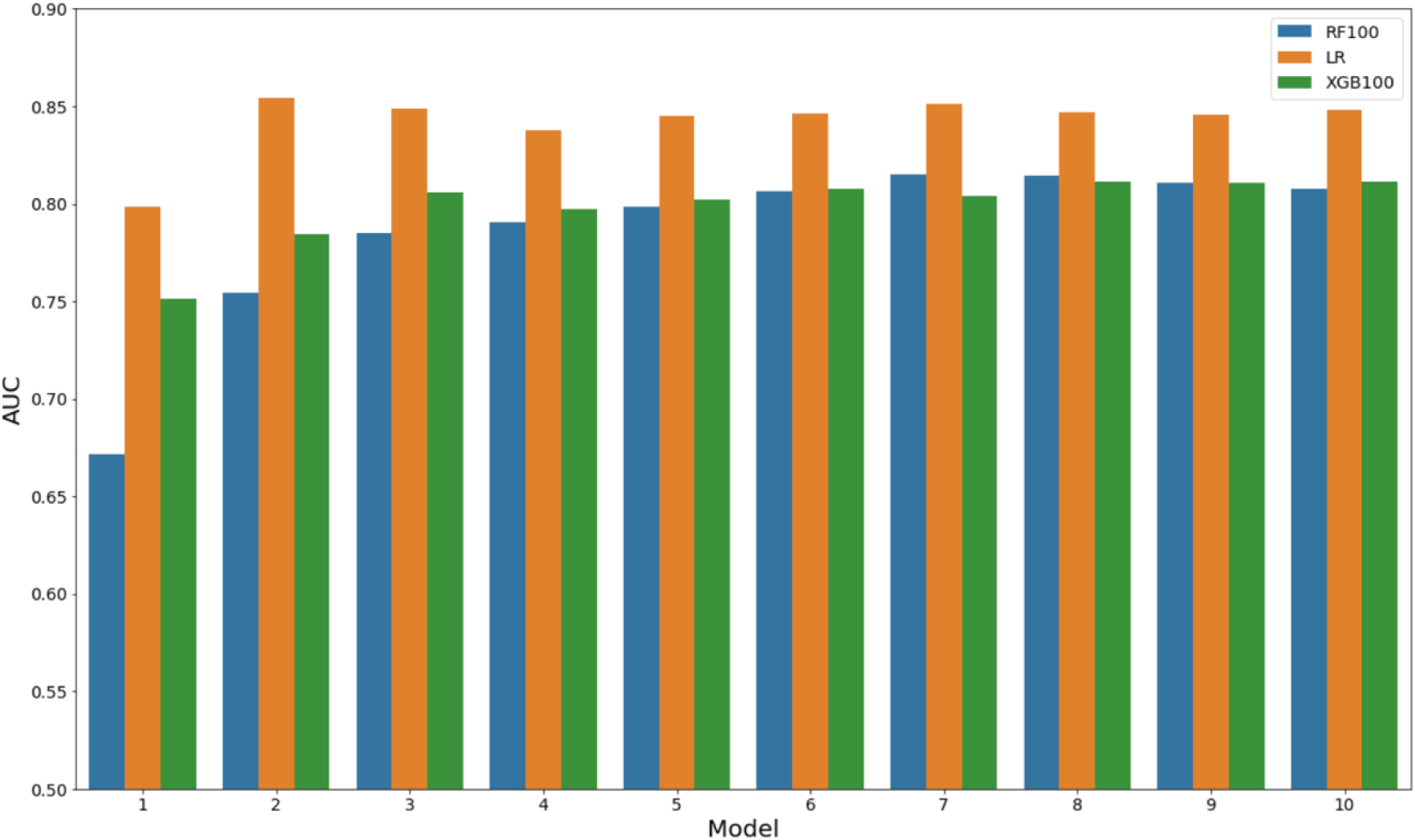
Performance comparisons of RF100, LR, and XGB100 models using top *k* = 1, 2, …, 10 features selected using MRMR_auc method.

### Machine learning based re-ranking of DEGs

Due to the small dataset and the instability of feature selection methods, the top seven DEGs selected in each fold might be different. Note that we conducted 10 runs of 10-fold cross-validation procedure. Thus, we chose seven DEGs 100 times to train and evaluate the LR model. To determine the importance of each gene, we assigned each gene a score indicating how many times (out of 100) this gene had been selected among the top seven genes used to train the classifier. Then, we simply normalized the scores by dividing by 100 such that gene importance scores of 1.0, 0.87, and 0.0 correspond to genes that have been selected 100, 87, and zero times, respectively. Additional file 1: Table S4 reports the gene importance scores for the 108 DEGs. Only 31 genes have importance score greater than zero. The top 15 genes and their importance scores are shown in Figure 3. We noted that three genes (DDIT4, RHAG, and AREG) had been consistently selected in each time.

**Fig. 3.**
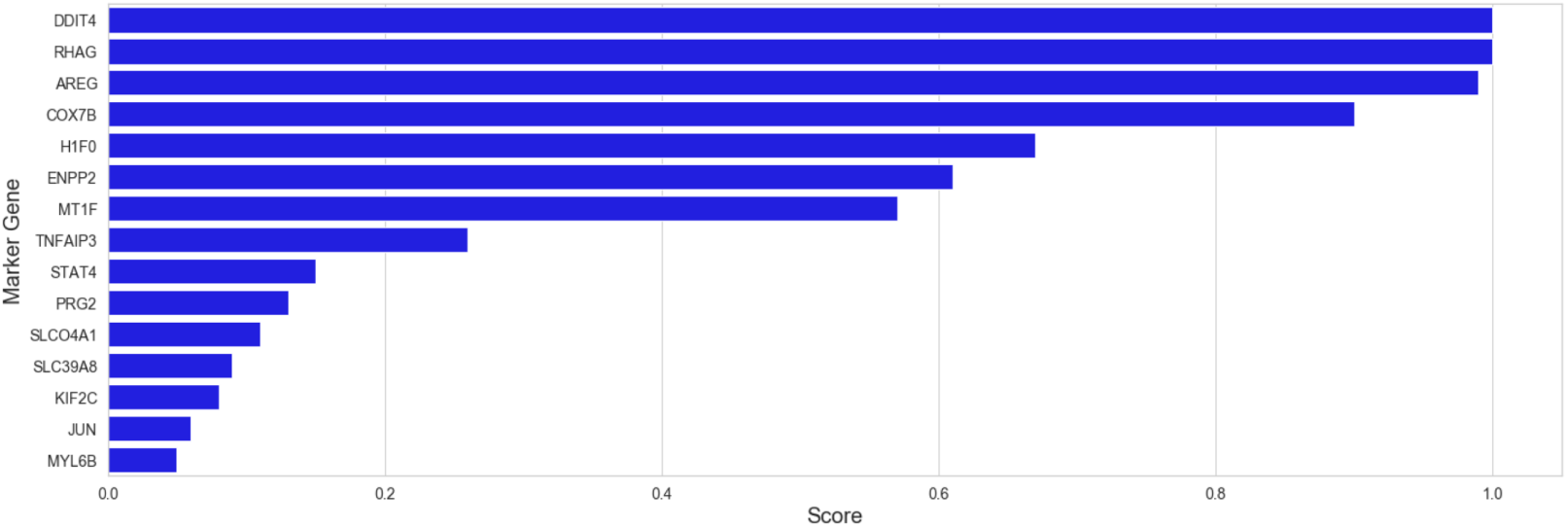
Top 15 gene markers identified using the proposed machine learning based DEGs re-ranking method.

In summary, our machine learning based refining of DEGs outcome reduced the number of DEGs from 108 to 31 and provided an alternative ranking of these genes. Next, we show how to use this ranking to determine the minimum set of DEGs that best discriminate between pediatric sepsis survivals and non-survivals.

### A 10-gene signature of mortality in pediatric sepsis

We used the top 15 genes in Figure 3 to search for a minimal set of genes that best discriminates between pediatric sepsis survivals and non-survivals. Specifically, for top *k* = {4, 5, …, 15} genes, we obtained the average ROC curves of LR models estimated using 10 runs of 10-fold cross-validation procedure (See Additional file 2: Figure S4). We found no improvement in the ROC curve when using more than top 10 genes. Figure 4 shows the boxplots of the normalized gene expressions of these 10 genes. Interestingly, all 10 genes are up-regulated. The most expressed genes are COX7B and DDIT4 while the least expressed genes are PRG2 and AREG.

**Fig. 4.**
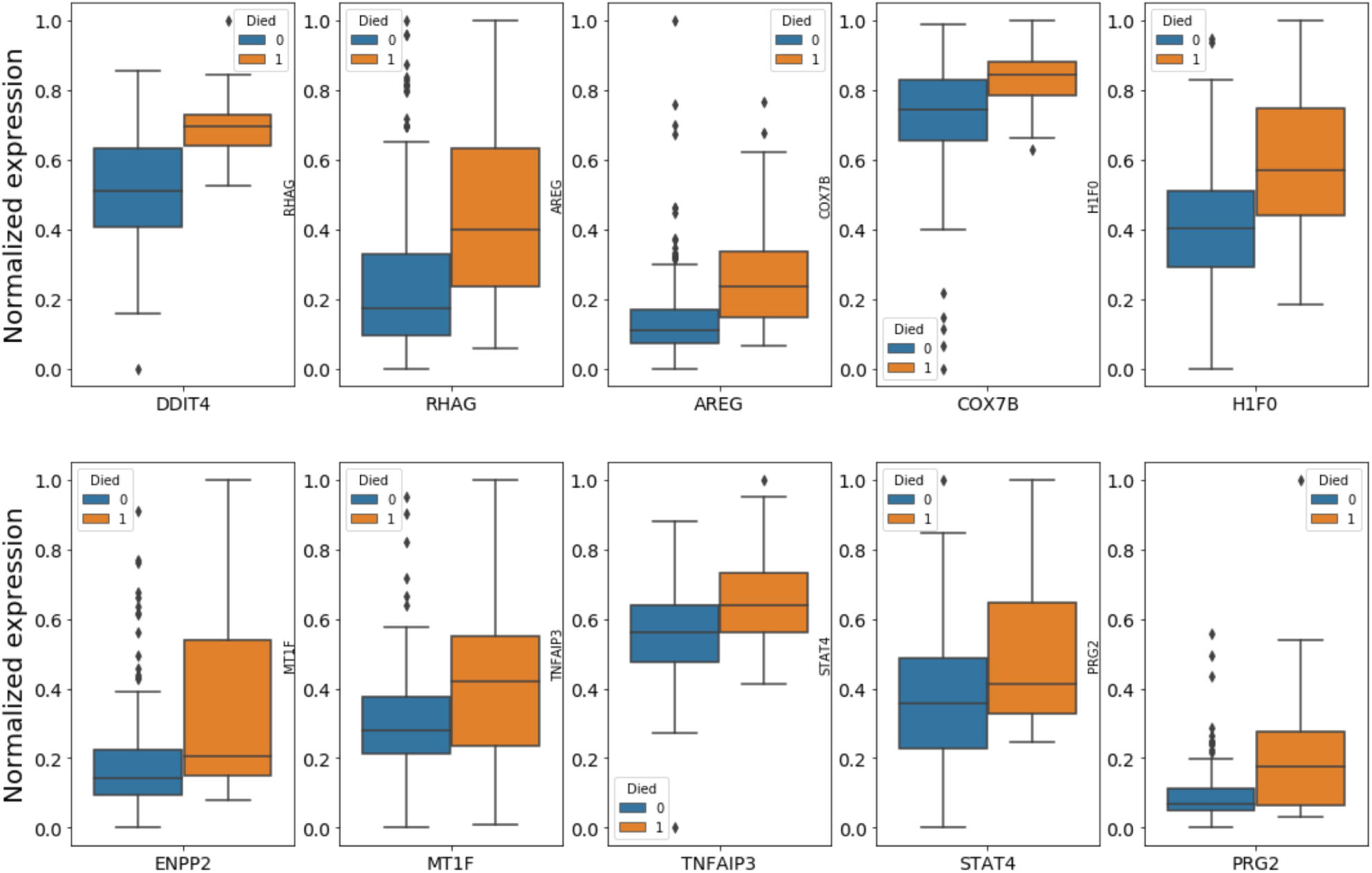
Boxplots for the normalized expressions of the 10 marker genes in survival and non-survival groups.

Using this panel of 10 marker genes, we compared the three machine learning algorithms considered in this study. We found that the ROC curve of the LR model almost dominates the two ROC curves for RF100 and XGB100 classifiers (Figure 5). Performance comparisons of these three classifiers are provided in Table 3. LR model has average AUC score of 0.89 while both RF100 and XGB100 have an average AUC score of 0.86. Moreover, the LR model has the best sensitivity, specificity, and MCC.

**Table 3:**
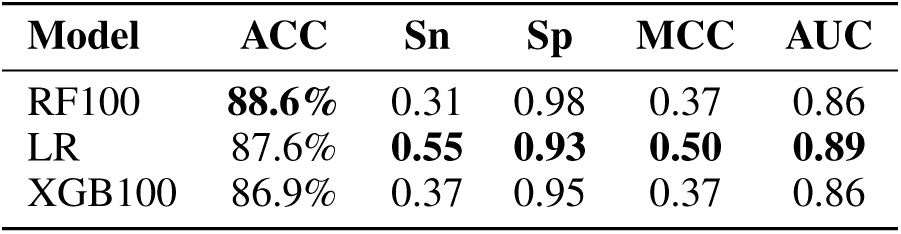
Performance estimates of different classifiers evaluated using 10 runs of 10-fold cross-validation procedure and top 10 gene markers.

| Model | ACC | Sn | Sp | MCC | AUC |
| --- | --- | --- | --- | --- | --- |
| RF100 | **88.6%** | 0.31 | 0.98 | 0.37 | 0.86 |
| LR | 87.6% | **0.55** | **0.93** | **0.50** | **0.89** |
| XGB100 | 86.9% | 0.37 | 0.95 | 0.37 | 0.86 |

**Fig. 5.**
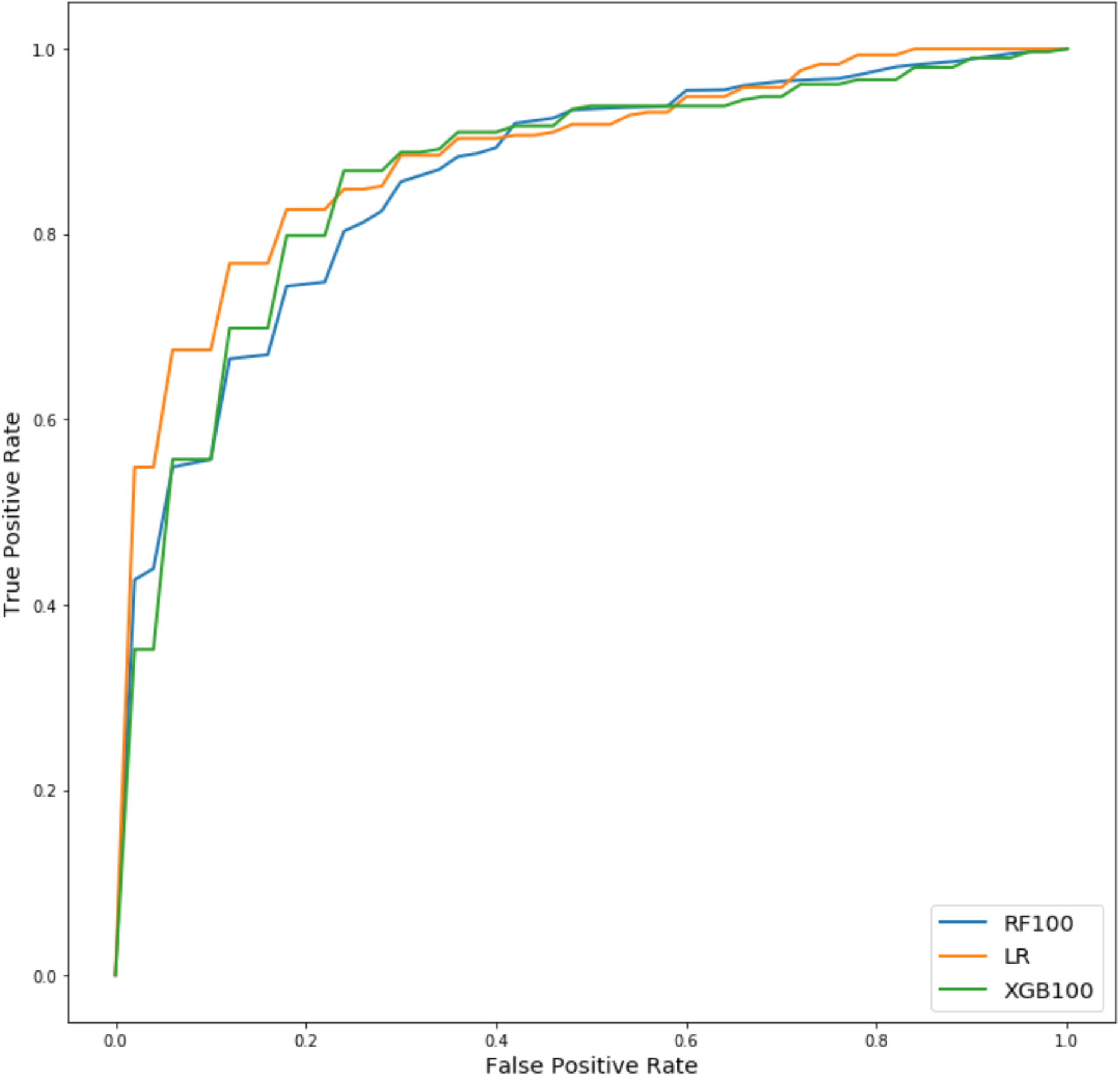
Average ROC curves of RF100, LR, and XGB100 models estimated using 10 runs of 10-fold cross-validation and 10 machine learning identified marker genes.

### Comparison of different gene ranking methods

We compared the LR model trained using the 108 DEGs to the LR models trained using only top 10 DEGs obtained using our proposed machine learning based gene ranking method (top10_ml) and two other ranking methods based on absolute fold change (top10_fc) and *p*-values (top10_pv). The average ROC curves of the four LR models are shown in Figure 6-a and the performance metrics of these models are reported in Table 4. The model using the 108 DEGs has the worst ROC curve and the lowest performance estimates. The model based on top 10 genes obtained using the absolute fold change ranking slightly outperformed the model based on top 10 genes ranked using the *p*-values. Finally, the model obtained using our proposed machine learning based ranking substantially outperformed all three models. Although all the models based on the three ranking methods had acceptable performance (i.e., AUC score ≥ 0.84), we found that the three sets of genes were not substantially overlapping with each other (See Figure 6-b). Every set of genes had at least 5 unique genes and the only common gene among the three sets was DDIT4. Figure 6 also visualizes the gene expression profiles for survival and non-survival patients in a 3D space defined by the top three marker genes in these three lists.

**Table 4:** Performance estimates of LR classifiers evaluated using 10 runs of 10-fold cross-validation procedure and different set of genes.

| Gene set | ACC | Sn | Sp | MCC | AUC |
| --- | --- | --- | --- | --- | --- |
| DEGs | 80.3% | 0.41 | 0.87 | 0.26 | 0.75 |
| top10_fc | 85.7% | 0.41 | 0.93 | 0.36 | 0.85 |
| top10_pv | 86.2% | 0.40 | **0.94** | 0.38 | 0.84 |
| top10_ml | **87.6%** | **0.55** | 0.93 | **0.50** | **0.89** |

**Fig. 6.**
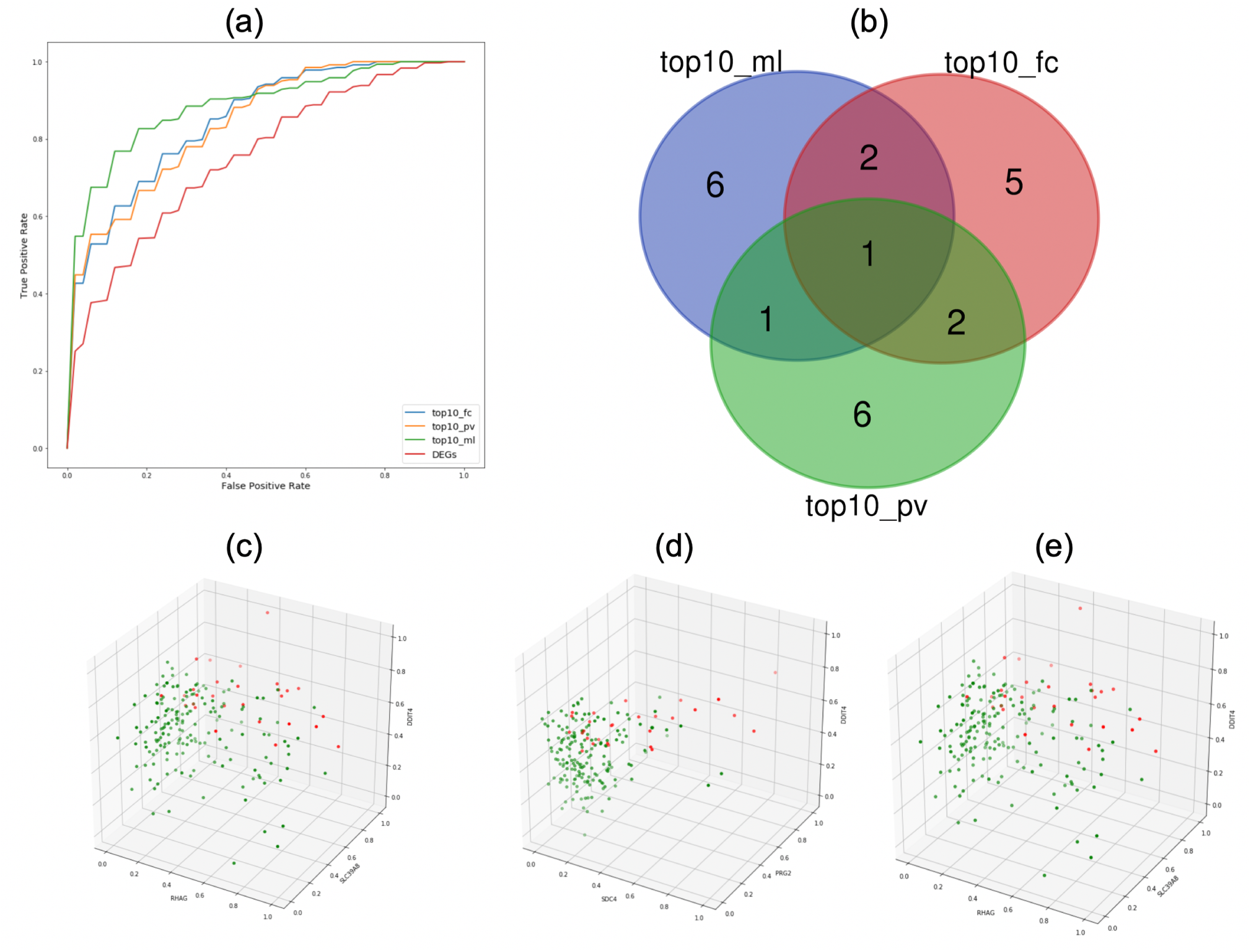
Comparisons of three gene ranking methods. (a) ROC curves of LR models evaluated using 108 DEGs and top 10 marker genes determined using fold change (top10_fc), *p*-value (top10_pv), and proposed machine learning method (top10_ml). (b) Venn diagram of the these three lists of 10 marker genes. Visualization of survival (green) and non-survival (red) samples in a three dimensional space based on the top three genes in (c) top10_fc, (d) top10_pv, and (e) top10_ml.

## Discussion

Differential expression (DE) analysis has been widely used to analyze gene expression profiles and uncover the underlying biological mechanisms for complex diseases [33, 34]. In general gene expression profiles are characterized with high dimensionality (tens of thousands of genes) and high pairwise correlations between genes. Therefore, the outcome of DE analysis tools often includes hundred(s) of highly correlated genes (see Additional file 2: Figure S2). Therefore, it is impractical to use all DEGs for developing diagnostic and prognostic prediction tools. In general, identifying a gene signature (a small set of marker genes) can be done using domain knowledge or data-driven approaches [14]. In this study, we presented a data-driven approach to prioritize the marker genes using an instance of the MRMR feature selection algorithm for selecting genes with the highest AUC for predicting the pediatric sepsis mortality and the minimal redundancy among selected genes in terms of Pearson’s correlation coefficients.

An interesting finding in our analysis is that the widely used performance metrics such as sensitivity, specificity, and AUC might not be sufficient to draw accurate conclusions regarding how different models compare to each other particularly when models are very competitive with each other and there is no model with a ROC curve that dominates the ROC curves for the remaining models. Another interesting finding is related to the observed surprisingly superior performance of LR models compared with RF100 and XGB100 models. This superior performance combined with the fact that LR models are linear interpretable models make LR algorithm a preferred choice for developing prediction models based on gene expression profiles as long as marker genes can be reliably identified.

It should be noted that supervised machine learning algorithms combined with feature selection methods could be directly applied to identify marker genes from the entire transcriptomic profiles. However, this approach suffers two major limitations. First, the computation time might be extremely long because some feature selection methods (including MRMR, feature selection based on genetic algorithms [35], and network-based feature selection [36]) have expensive computational time proportion to the number of features. Second, it is challenging to apply functional enrichment analysis to the identified set of marker genes because of the small number of identified genes and the lack of significant redundancy among these genes [19]. Therefore, it is less likely that these genes share any common functional pathways. The present approach utilizes supervised feature selection to refine the outcome of statistical DE analysis. It will be interesting to explore novel approaches for separately applying statistical DE and supervised feature selection to entire gene expression profiles and then integrate the outcome of the two methods. For example, NetworkAnalyst tool supports comprehensive meta-analysis of multiple gene lists through heatmaps, Venn diagrams, and enrichment networks. One interesting way for obtaining more than one list of DEGs is to obtain them using different statistical and machine learning approaches.

Our DE and machine learning analyses suggested three 10-gene marker lists for predicting mortality in pediatric sepsis with average AUC score ≥ 0.86. These three lists had only one gene in common, which suggests the existence of multiple data-driven gene signatures for mortality in pediatric sepsis. Similar observation had been reported by Sweeney et al. [19] where the authors had reported four sets of sepsis marker genes with only few genes in common. This underscores the need for independent validation set as well as wet laboratory experiments to validate some of these markers and confirm the reported biological insights.

## Conclusions

We have identified a signature of 10 marker genes for reliably predicting mortality in pediatric sepsis. These 10 genes have been determined using a novel machine learning data-driven approach for re-ranking and selecting an optimal subset of 108 DEGs identified via a secondary analysis of, to the best of our knowledge, the largest publicly available transcriptomic cohort study for pediatric sepsis. Our on-going work aims at: i) validating our proposed 10-gene signature using an independent test set; ii) testing and evaluating the proposed approach for identifying reliable biomarkers for challenging biomarker discovery tasks in critical care settings such as diagnosing and endotyping sepsis and Acute Respiratory Distress Syndrome (ARDS); iii) Adapting our approach for single cell gene expression analysis [38, 39].

## Supplementary information

**Additional file 1:** Supplementary Tables S1-S4

**Additional file 2:** Supplementary Figures S1-S4

