## Additional File 2 for "Machine Learning Based Refined Differential Gene Expression Analysis of Pediatric Sepsis"

**Fig. S1** Volcano plot of gene expression genes in pediatric sepsis. Genes highlighted in blue represent significantly up-regulated (right) and down-regulated (left) genes.

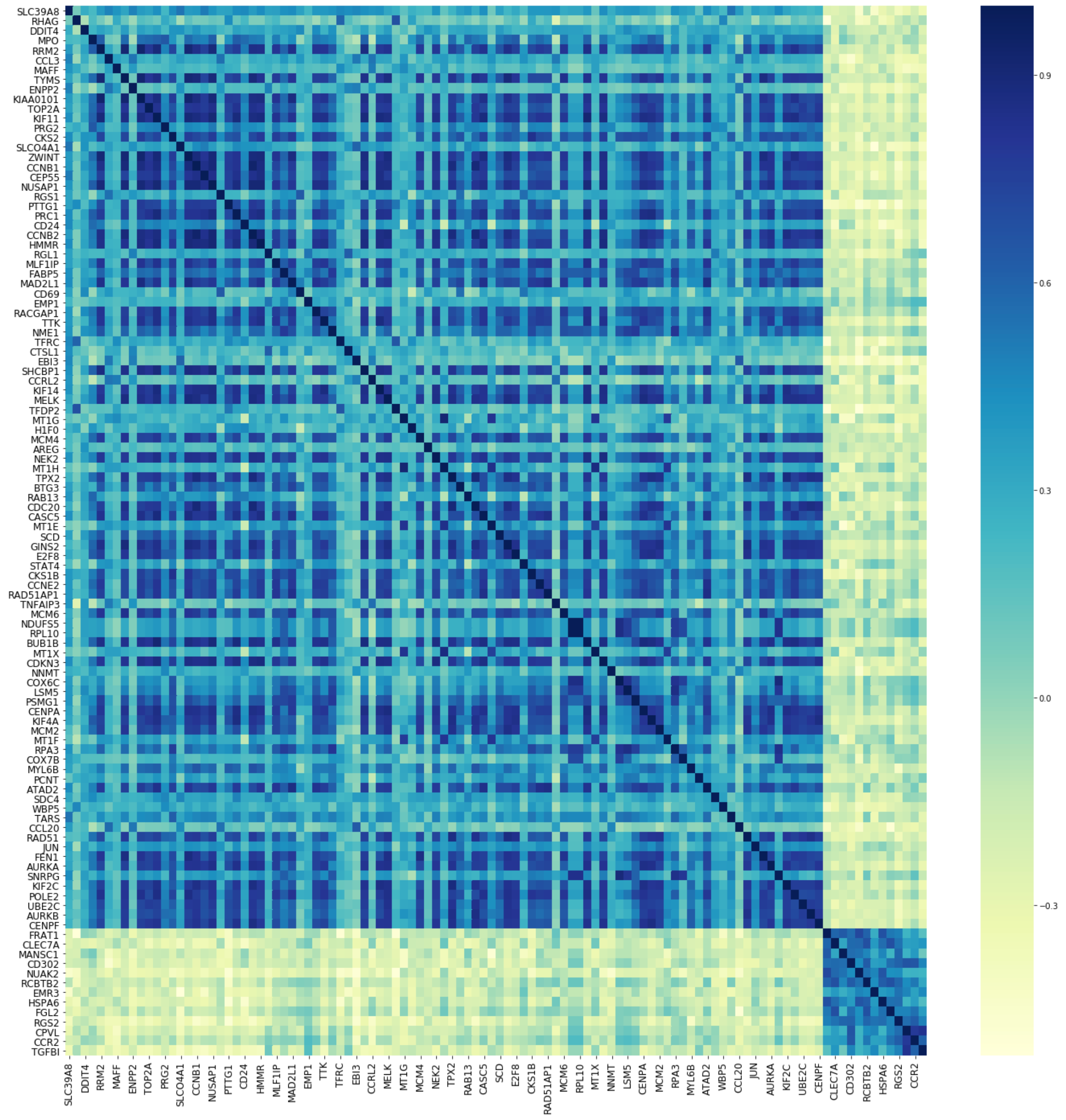

**Fig. S2** Heatmap of Pearson’s correlation coefficient matrix of the 108 DEGs.

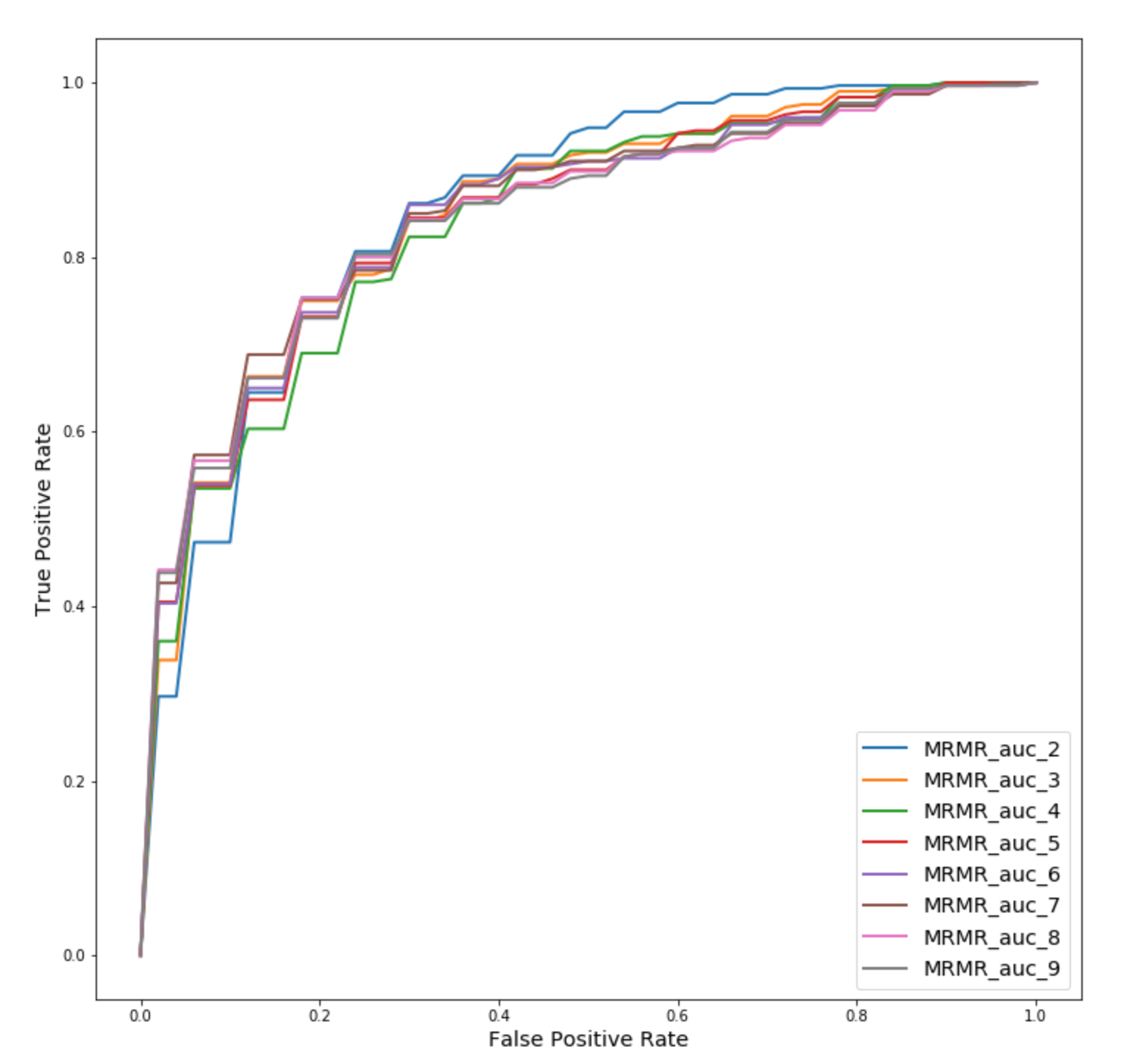

**Fig. S3** Average ROC curves of LR models estimated using 10 runs of 10-fold cross-validation and MRMR_auc for selecting top 2,3,…,9 marker genes from the 108 DEGs.

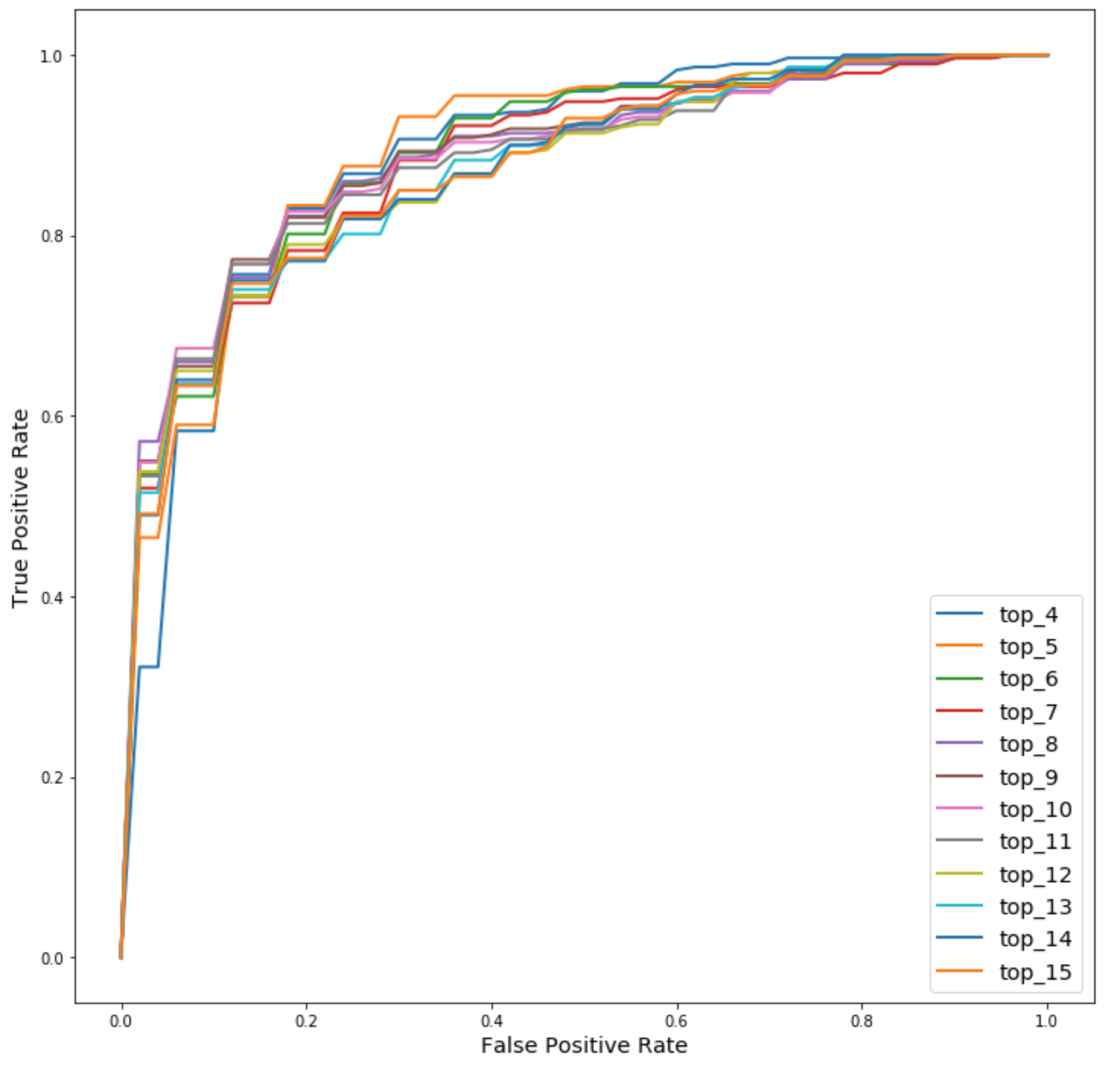

**Fig. S4** Average ROC curves of LR models estimated using 10 runs of 10-fold cross-validation and top re-ranked 4,5,…15 marker genes.
